# Dynein collective behavior in mitotic nuclear positioning of *Saccharomyces cerevisiae*

**DOI:** 10.1101/2020.06.23.166769

**Authors:** Kunalika Jain, Neha Khetan, Saravanan Palani, Chaitanya A. Athale

## Abstract

Positioning the nucleus at the bud-neck prior during *Saccharomyces cerevisiae* mitosis during anaphase involves pulling forces of cytoplasmic dynein localized in the daughter cell. While genetic analysis has revealed a complex network positioning the nucleus, quantification of the forces acting on the nucleus and dyneins numbers driving the process has remained difficult. In order to better understand the role of motor-microtubule mechanics during nuclear positioning and the role of dynein, we have used a computational model of nuclear mobility in *S. cerevisiae* and reconciled it to the mobility of labelled spindle pole bodies (SPBs) measured by quantifying fluorescence microscopy time-series. We model the apparent random-walk mobility of SPBs by combining diffusion of the nucleus and active pushing of MTs at the cell membrane. By minimizing the deviation between tracks of fluorescently tagged SPBs and simulations, we estimate the effective cytoplasmic viscosity to be 0.5 Pa s. The directed transport of nuclei during the budding process is similarly quantified by tracking the daughter SPB (SPB-D) in experiment. Using force-balance, we find 2 to 8 motors are required to pull the nucleus to the bud-neck. Simulations of the cytoplasmic MT (cMT) ‘search and capture’ by dynein suggest single motor binding is followed by a rapid saturation of number of bound motors. The short time and length of MT interactions with the cortex and minimal collective dynein force required, predict a functional role for dynein clustering in nuclear positioning.

## Introduction

Centrosomes nucleate radial array of microtubule (MT) arrays or asters that are vital for mitotic spindles and maintaining cell geometry in interphase in animal cells. A structure comparable to centrosomes, the spindle pole body (SPB), has been identified in the model budding yeast *Saccharomyces cerevisiae* (Seybold and Schiebel, 2013) consisting of centrins and *γ*-tubulin (Schiebel and Bornens, 1995) and regulating nuclear and cytoplasmic MT nucleation and number (Knop et al., 1999). During mitosis astral or cytoplasmic MTs (cMTs) play a vital role in positioning the nucleus by asymmetric pulling forces, but unlike in higher eukaryotes the asymmetry appears to be generated by the differential transport of force generators like dynein into the daughter cell (Markus et al., 2012a; Chen et al., 2019). Once localized, dynein interacts with the astral MTs and position the spindle at the onset of anaphase during *S. cerevisiae* mitosis (Carminati and Stearns, 1997; Adames and Cooper, 2000; Baumgärtner and Tolić, 2014). Evidence has accumulated for an ‘offloading model’ of dynein transport in an inactive form to the plus ends of astral microtubules (MTs) before it binds the cortex (Lee et al., 2005; Markus and Lee, 2011). In turn a protein She1 inhibits dynein movement selectively in the mother cell by resulting in asymmetric pulling forces into the daughter cell (Markus et al., 2012b). In the daughter cell, cortical dynein both walks on MTs and destabilizes them (Estrem et al., 2017). This role of cortical dyneins pulling on cMTs to position spindles has also been well studied in the *C. elegans* embryos (Grill et al., 2001) and human cells (Kiyomitsu and Cheeseman, 2013), while in *Schizosaccharomyces pombe*, the combined effect of MT lengths and motor-numbers result in nuclear oscillations (Svoboda et al., 1995; Ding et al., 1998; Yamamoto et al., 2001). The orientation of spindles in animal cells has been shown to be mechanically regulated by forces acting through the number and length of astral MTs interacting with dyneins localized at the cell cortex (O’Connell and Wang, 2000; Grill et al., 2001; Kotak et al., 2012). Models proposed for spindle positioning suggest MTs interact with cortical dyneins either (i) end-on (Kozlowski et al., 2007) or (ii) laterally (Adames and Cooper, 2000). Indeed, it appears a end-on interactions transition to lateral ones, and might be necessary for multiple motors to act on astral MTs and generate sufficient force to position a spindle or nucleus. Together, these form an increasingly complex picture of dynein and cMTs in spindle positioning during budding yeast mitosis. Despite the progress in imaging dynamics of the process (Lammers and Markus, 2015; Fees et al., 2017), quantification of the forces driving nuclear positioning remains incomplete. Therefore there exists a need for a detailed mechanical model reconciled to experiments.

The *S. cerevisiae* dynein has become a model for understanding dyneins generally, due to advances from single-molecule dynamics demonstrating a high degree of processivity (Reck-Peterson et al., 2006) that has been attributed to tension-dependence of coordinated stepping of pairs of heads (Qiu et al., 2012). A clear picture of single molecule coordination has been emerging from the demonstration of force-dependent bidirectional stepping of dynein (Gennerich et al., 2007) and load-sharing between heads (Belyy et al., 2014), and most interestingly an asymmetry in detachment of the motor from MTs depending on the direction of the load force (Nicholas et al., 2015). The potential connection between these single-molecule properties and collective transport of MTs was recently demonstrated in our study combining experiment and simulations, with ensembles in excess of 8 to 10 motors resulting in cooperative transport, explained by the asymmetry in load dependent detachment rates of the single motor (Jain et al., 2019). However, the implications of these *in vitro* and *in silico* observations for the precise number of dynein molecules *in vivo* remains unclear. Thus a mathematical model reconciled to experimental data from *S. cerevisiae* could provide a clearer picture of the nature and magnitude of the forces involved.

The mobility of organelles through the cytoplasm is expected to be dominated by viscosity, due to the low Reynolds number of organelle movement. Estimations of viscosity of budding yeast cells have ranged from 0.3 to 0.7 Pa s (Baumgärtner and Tolić, 2014). In contrast, the value of the fluid phase viscosity using electron spin resonance (ESR) spectroscopy in the fungus *Talaromyces macrosporus* ranged from 2.5 to 15 × 10^−3^ Pa s, while in *Penicillium sp*. ranged between 3.5 to 4.8 × 10^−3^ Pa s, depending on the developmental stage (Dijksterhuis et al., 2007). In addition, the estimation of intracellular viscosity in cell culture cells typically estimates microviscosity or fluid phase viscosity, which for cytoplasm is reported to be only 1.2 to 1.4 × 10^−3^ Pa s (Seksek et al., 1997; Fushimi and Verkman, 1991). However, larger objects such as organelles or micron-scale particles that encounter intracellular structures and other organelles could experience a higher effective viscosity, which can even reach values as high as 2 Pa s (Bacher et al., 2004). This object size dependence of cytoplasmic effective viscosity has been attributed to a ‘sieving’ effect in the cytoplasm (Seksek et al., 1997; Mika et al., 2010). It also implies the a scale-specific method of estimation of the drag force acting on nuclei will be required.

MTs nucleated from centrosomes or spindle pole bodies (SPBs) that do not form part of the spindle are referred to as astral or cytoplasmic MTs (cMTs). They contact the cell cortex, experience pulling forces, and determine the division plane. The first stage of this is reminiscent of ‘search and capture’ of dynamic MTs seen during spindle assembly (Holy and Leibler, 1994). In large cells additional directional cues such as gradients of regulatory proteins that affect MT growth dynamics in *Xenopus* oocyte meiosis II (Athale et al., 2008, 2014) or motor gradients in mouse oocyte meiosis I (Khetan and Athale, 2016) are also thought to play a role. Once captured, cortical motors generate pulling forces on cMTs as seen in early embryonic divisions of *Caenorhabditis elegans* (Kozlowski et al., 2007; Grill et al., 2001). In yeast cells, in addition to these cortical interactions the ‘capture’ of MTs by motors is also thought to be followed by their depolymerization (Adames and Cooper, 2000). Such depolymerization while walking on the MTs could compensate for an important source of mechanics in spindle positioning based on MT pushing by polymerization seen *in vitro* (Dogterom et al., 2005; Dogterom and Yurke, 1997). The most recent modeling study of nuclear positioning in yeast based on MT-motor mechanics is limited by the fact of continuum assumptions about motors and MTs (Sutradhar et al., 2015), while the number of motors and MTs seen in experiment are small in number, and likely to be subject to fluctuations. This suggests the a model that takes into account individual MTs and single motor mechanics is required for an improved understanding of the biophysical basis of nuclear positioning in *S. cerevisiae* mitosis.

Here, we have used a video microscopy approach to analyze time-series of *S. cerevisiae* nuclei by single particle tracking of fluorescently labelled spindle pole bodies (SPBs). The mobility of pre-budding cells appears to be random, with both diffusion and MT pushing playing a role. We develop a model nuclear mobility driven by dynamic MTs pushing on the cell membrane and diffusion and estimate the effective cytoplasmic viscosity, by reconciling theory with experiment. Based on this quantification, we then examine intracellular SPBs mobility during budding, as the nucleus is pulled into the bud neck. We model the process using a minimal geometry and based on force balance and direct simulations (Data S2), estimate the collective force driving nuclear positioning, by comparison to experiment.

## Materials and Methods

### Yeast growth conditions and live imaging

Yeast strains expressing either a GFP-tagged copy of the SPB protein Spc42 or a dual-labelled strain with Spc42 tagged with mCherry and GFP-Tub1 (Caydasi and Pereira, 2009) were used (Table 1). Yeast growth methods and growth media used have been described in a previous study (Sherman, 1991). Log phase cells were cultured in filter-sterilized yeast peptone dextrose medium containing 0.1 mg/l adenine (YPDA) medium at 30°C for time-lapse imaging. Time-lapse movies were acquired for 2 to 3 hours at 30°C under temperature controlled incubation chamber and yeast cells were immobilized on CellASIC microfluidic yeast plates (Y04C size) with a constant flow of YPDA medium for 3 hours. Time-lapse movies were acquired using the spinning disk confocal microscope (Andor Revolution XD imaging system, equipped with a 100x oil immersion 1.45 NA Nikon Plan Apo lambda objective (Nikon Corp., Japan), a Confocal Yokogawa CSU-X1 unit, an Andor sCMOS ZYLA detector, and Andor iQ software (Oxford Instruments, UK). Z-stacks comprised of fifteen 0.5 *µ*m spaced slices were generated for GFP-Tub1 and Spc42-3xmCherry at 1 minute intervals.

**Table 1:**
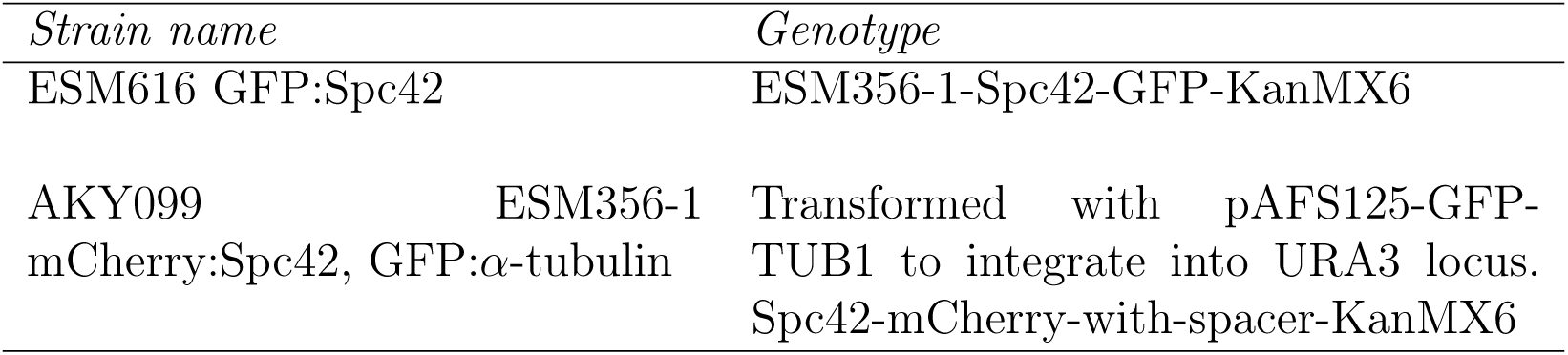
Strain list: The list of yeast strains used in this study. These strains have been described in a previous study by Caydasi and Pereira (Caydasi and Pereira, 2009).

### Image analysis

Image time series of yeast cells were contrast adjusted, denoised using a median filter (radius-1 pixel), and histogram normalized using Image J ver 1.45s (Schneider et al., 2012) with the *Stackreg* plugin to align stacks of images that underwent drift during acquisition. The channel with the SPB is labelled by Spc42-3xmcherry. The signal in the mCherry channel was tracked using a nanometer precision filament and particle tracking program FIESTA (Ruhnow et al., 2011) implemented in MATLAB R2020a (Mathworks Inc., MA, USA), with a threshold search velocity of 50 nm/s and a minimal framepersistence of 15 minutes The XY co-ordinates of the tracks were plotted and overlaid on the DIC channel used to visualized and categorize cells into either pre-budding or budding stages. Further, in budding stages the analysis was considered with cells were SPB movement to mother and bud cells was unambiguous with track length ≥ 50 time frames.

### Simulations

2D simulations were performed using Cytosim, a C++ based Langevin dynamics simulation engine (Nedelec and Foethke, 2007). Cell geometry was implemented as an ellipse with a circular nucleus initialized at the cell-center. The SPBs are embedded on the nuclear envelope such that their relative movement along the surface is restricted as the statistics in experiments is calculated prior to SPB segregation (pre-bud) and prior to spindle elongation after spindle formation (for the budding cells). Each SPB nucleates two MTs that undergo dynamic instability modeled by the two state-four parameter model (Verde et al., 1992). Dynamic instability parameters are taken from primarily two papers that quantify dynamics in a cell-cycle stage specific manner (Table 2). The system components are confined in the cellular space. The dynein motors are localized at the cortex along the arc of the ellipse as chosen arbitrarily to mimic the curvature of the bud. The integration time was chosen to be smaller than the fastest time-scale (Table 3). In the pre-budding cells, the cytoplasmic viscosity was varied and the difference in mean simulation outputs compared to the same metric from experimental quantification were rank sorted, based on the minimization of deviation between experiment and simulations. This validation from experiments was used to estimate the optimal effective cytoplasmic viscosity *η*_*cyto*_, that was then used for budding calculations. A typical simulations run required approximately 20 minutes on a 12 core Intel Xeon machine with 16 GB RAM for total time 20 minutes, time-step Δ*t* of 10^−3^ seconds and a configuration involving 1 nucleus with a pair of SPBs nucleating a pair of MTs and up to 200 motors.

**Table 2:** MT dynamic instability parameters used in simulations of both pre-budding and budding cells.

| Cell stage | $v_g$ | $v_s$ | $f_c$ | $f_r$ | Reference |
| --- | --- | --- | --- | --- | --- |
| *Pre-bud* | 0.009 $\mu\text{m/s}$ | 0.03 ($\mu\text{m/s}$) | 0.007 $\text{s}^{-1}$ | 0 $\text{s}^{-1}$ | (Carminati and Stearns, 1997) |
| *Pre-anaphase* | 0.03 $\mu\text{m/s}$ | 0.05 ($\mu\text{m/s}$) | 0.023 $\text{s}^{-1}$ | 0.03 $\text{s}^{-1}$ | (Fees et al., 2017) |

**Table 3:** Simulation parameters determining cell geometry and viscosity.

| *Symbol* | *Description* | *Stage* | *Value* |
| --- | --- | --- | --- |
| $d_{nucl}$ | Nuclear diameter | Pre-budded | 1 $\mu\text{m}$ |
| - | - | Budded | 1 $\mu\text{m}$ |
| $d_{cell}$ | Cell diameter along long and short axis | Pre-budded | 6, 5 $\mu\text{m}$ |
| | - | Budded | 8, 4.5 $\mu\text{m}$ |
| $n_{SPB}$ | Number of SPBs per cell | Pre-budded and Budded | 2 |
| $n_{MT}$ | Number of MTs per SPB | Pre-budded and Budded | 2 |
| $\eta_{cyto}$ | Yeast cytoplasmic viscosity | Pre-budded and Budded | Varied from $10^{-2}$ to $10^1$ Pa.s |
| $\theta_{spb}$ | Angle between the SPBs | Pre-budded | $0^\circ$ |
| | | Budded | $180^\circ$ |

### MSD and diffusivity analysis of trajectories

We have used a standard method of MSD calculations as previously described (Arcizet et al., 2008; Athale et al., 2014; Khetan and Athale, 2016) based on the 2D positional coordinate of the filament (**r**) using a sliding-window approach of increasing time intervals (*δ*t) based on the expression:

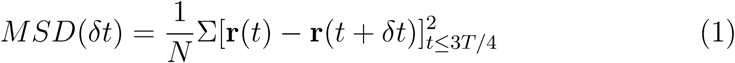

where *N* = *T/δt*, the number of steps of a given time interval, *t* is the time point and *T* is the total time. The time-series was analyzed only for values of *δt* less then or equal to 0.75 × *T*, based on previous reports (Arcizet et al., 2008; Michalet, 2010; Jain et al., 2019), and MSD was plotted as a function of *δt*.

We fit two models of effective diffusion to the MSD data from both experiment and simulation: *(a)* anomalous diffusion and *(b)* diffusion with drift. The *(a)* anomalous diffusion model is described by the expression:

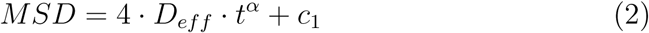

where *D*_*eff*_ is the effective diffusion coefficient, *t* is time, *α* is the parameter of anomalous diffusion and *c*_1_ is the error in detection. The *(b)* diffusion with drift model is given by:

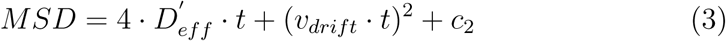

where 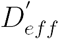 is the alternative estimate of the effective diffusion coefficient, *t* is time, *v*_*drift*_ is the drift velocity and *c*_2_ is the error in detection.

## Results and Discussion

### Diffusive mobility of spindle pole body (SPB) in prebudding cells

We have followed the dynamics of nuclear positioning in dividing *S. cerevisiae* mitosis by video microscopy, using a strain with a fluorescently tagged SPB (Spc42-3xmCherry) and GFP labeled tubulin, described previously (Caydasi and Pereira, 2009). Based on the morphology of the cell in the DIC image, the pre-budding stage is selected and single particle tracking in the mCherry channel is used to follow SPB mobility as seen in the representative Video SV1 and Fig. 1A). The XY trajectories of up to 20 SPBs normalized to their initial position appear to explore the cytoplasmic space performing an apparent random walk (Fig. 1B).

**Figure 1:**
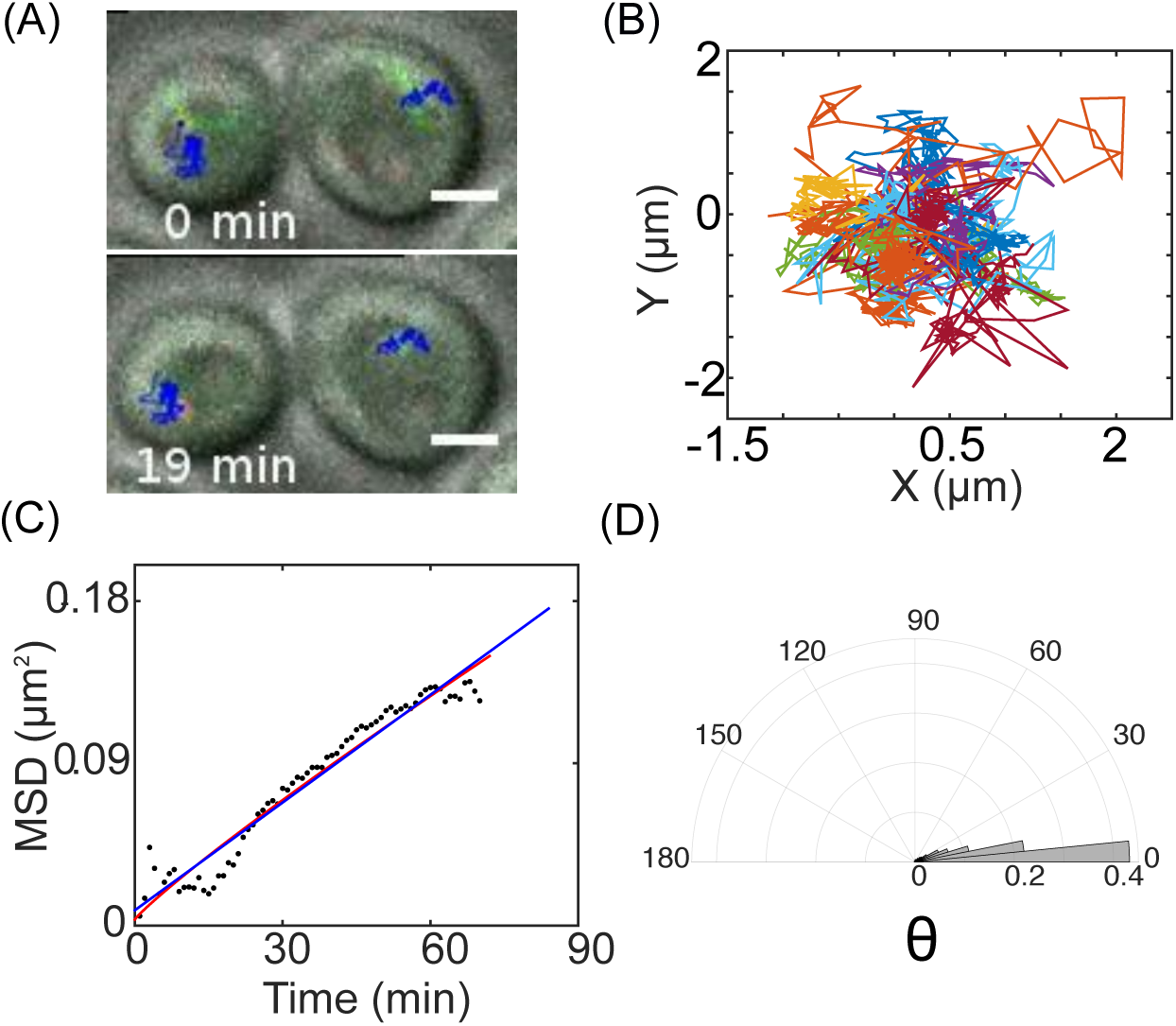
Tracking SPB mobility in the pre-budding (early G1) stage. (A) Representative image of a time series of yeast cells selected based on morphology in DIC in the pre-budding stage (gray) overlaid with the Spc42-3xmCherry (red) and GFP-Tub1 (green) channels (corresponding to Video SV1). The time points 0 min and 19 min correspond to start and end of the stack. Tracks of SPB mobility(blue) are overlaid. Scale bar 2 *µ*m. (B) Multiple SPB tracks (n = 20) are plotted with their origin normalized showing apparent random movement. (C) The MSD (*µ*m^2^) for a representative pre-budding SPB is plotted as a function of time (min) and (D) the representative angular histogram of the angle between subsequent time points of SPB position subtended at the origin are plotted.

We estimated the MSD values for increasing time intervals from experimental trajectories, as described in the methods section, and fit MSD data to a model of anomalous diffusion (Equation 2). The resulting fit value of effective diffusion coefficient *D*_*eff*_ was 0.0565 ± 0.09 *µ*m^2^/min and of the coefficient of anomaly of diffusion *α* was 1.172±1.3 (mean±s.d.) for SPBs in cells not yet budded (Figure 1C). The angular frequency distribution of the angle formed between subsequent positions of the SPB show a relatively tight distribution (Figure 1D). The value of *α* ∼ 1 is indicative of diffusive mobility. On the other hand the estimate of *D*_*eff*_ is affected by a combination of thermal random motion as well as pushing of dynamic MTs at the cell boundary.

In previous work, drag forces have been successfully estimated by measuring the viscosity of the cytoplasm by microrheology of vesicle (Hayashi et al., 2013) and lipid granule mobility (Grill et al., 2001). However, nuclear mobility determined not only by thermal forces opposed to drag, but also forces due to MT-pushing at the boundary. In order to account for both effects, we use a numerical simulation of the factors affecting nuclear mobility and compare the statistics to experimental measures.

### Estimating cytoplasmic viscosity by fitting a nuclear mobility model to experiment

We aimed to find the effective viscosity of the cytoplasm from simulations, by simulating the well parameterized model of MT mechanics and diffusion in Cytosim (Nedelec and Foethke, 2007), and minimizing the variation from SPB mobility statistics from experiment. The nucleus is modeled with a pair of SPBs embedded in the periphery, nucleating dynamic MTs based on the factors driving pre-budding nuclear mobility (Figure 2A). The parameters of cell and nuclear sizes, MT dynamics and flexural rigidity of the MTs are all taken from literature (Table 3). Simulations were run for increasing cytoplasmic viscosity (*η*) from 10^−2^ to 10^1^ Pa s with the nucleus initialized at the cell center and a random orientation of the SPBs and MTs. Given the rigid nature of the cell boundary, growing MTs that contact the boundary produce an inward force of polymerization acting on the SPBs (Video SV2). Rotational diffusion of the nucleus results in a random repositioning of SPBs and their corresponding MTs (Figure 2B). Combined with stochasticity of length fluctuations, the pushing forces experienced by the nucleus are random both in magnitude and direction. The resultant trajectories of SPB positions transition from a restricted random walk, through effective diffusion to diffusion with drift (Figure 2C). Mobility of the SPBs for a low value of viscosity appears to be diffusion-dominated, while high values are dominated by MT-pushing. The MSD averaged over the ensemble of SPB trajectories also confirms the restricted diffusion at low viscosity, with SPBs exploring the cellular space almost completely for *η* of 0.5 Pa s with a ‘corralling’ effect seen due to the cell boundary (Figure 2D). The ‘corralling’ radius is ≈ 2 *µ*m, corresponding to radius along the short axis of the cell (Table 3). We find the effective diffusion coefficient *D*_*eff*_ decreases expectedly with increasing viscosity, while the anomaly coefficient *α* increases (Figure S1). The increase in *α* with *η* suggests the nuclear mobility in the low viscosity regime is dominated by thermal energy, while MT pushing dominates in the high viscosity, an effect also also reflected in the directionality *χ* measure. The SPB velocity however shows a consistent decrease suggesting diffusion dominated mobility is faster for exploring cellular space than MT pushing. The angle between subsequent positions of the SPB acts as a measure of rotational diffusion of the nucleus (Figure 2F). The values of *η* were varied and the deviation between simulation and experiment (Supporting Data S1) for each of the measured variables *D*_*eff*_, *α, χ*, velocity and *θ* were ranked. The ranks across these measures were summed as a function of *η* (Figure 2F), arrive at an estimate of 0.5 Pa s as the best match for experiments and taken to be the cytoplasmic viscosity *η*_*cyto*_.

**Figure 2:**
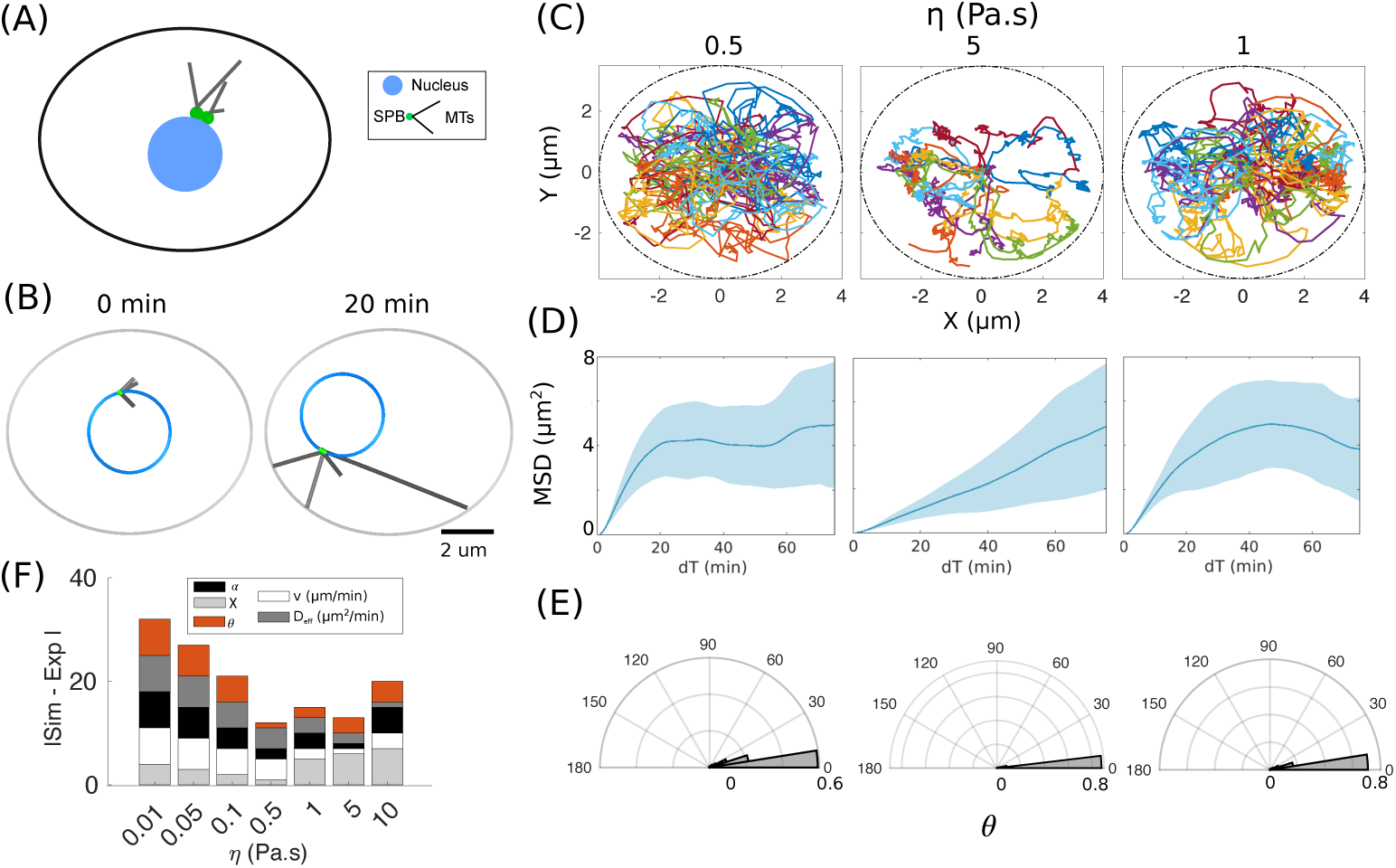
Estimating cellular viscosity from simulations of prebudding nuclear mobility. (A) Schematic of the pre-budding yeast set up in simulations with a nucleus (blue) with a pair of closely spaced SPBs (green) embedded on the nuclear periphery and two MTs nucleated per SPB. (B) Simulation outputs from time points 0 and 120 minutes (corresponding to Video SV2). Scale bar 2 *µ*m. (C-E) The viscosity (*η*) was varied to examine the effect on (C) XY trajectories of individual SPBs (n = 50), (D) MSD averaged over all SPBs (blue) with ± s.d. (light blue area) was fit to the anomalous diffusion model (Equation 2), to estimate the effective diffusion coefficient *D*_*eff*_ (*µ*m^2^/min) and anomaly coefficient *α* and (D) the frequency distribution of the angle *θ* made by subsequent positions of the SPB. (E) The bar graph depicts the deviation between the experiment and simulation as a function of *η* using a ranked difference minimization approach. The variables used were for the measures *α* (black), *D*_*eff*_ (dark-grey), *χ* (light-grey), *θ* (red) and velocity v (white). Simulation time = 100 minutes, N_*runs*_ = 50.

The cytoplasmic viscosity serves to damp random motion of the nucleus in the pre-budding stages, but during mitosis when the nucleus is transported to the bud-neck, it is expected to produce an opposing force due to drag. In order to understand the mechanics of the process, we proceed to quantify the *in vivo* mobility of the nucleus during the budding process, by tracking the SPB.

### Quantifying *in vivo* SPB motility during budding

During mitosis, the SPBs are duplicated and move to the diametrically opposite poles of the nucleus (Knop et al., 1997; Seybold and Schiebel, 2013). Based on their eventual fates those that enter the mother and daughter cells are referred to as SPB-M and SPB-D respectively. The time-series of punctate localization of Spc42-mCherry that mark the SPBs were tracked in cells that are budding as seen in the DIC channel (Video SV3(a)). We track the SPB-M and SPB-D mobility typically over 15 to 20 minutes (Figure 3A,D) and find mobility appears to be qualitatively more directional than pre-budding cells (Figure 3B,E). This is confirmed by quantification of the MSD based on the XY trajectories fit to the anomalous diffusion Equation 2 result in anomaly exponent *α >* 1.5 (Figure 3C,F, Table 4). Additionally, the SPB-D appears to migrate with a mean drift velocity of 17 nm/min while the SPB-M drift velocity is 4.1 nm/min based on fitting the MSD data to the diffusion and drift model (Equation 3). The heterogeneity in fit values could be the result from time-dependence of the processes driving SPB mobility. Transitioning of aster motility from random to directed in case of centrosome nucleated asters in *Xenopus* oocyte extracts involve a gradient of MT stabilization that increases length asymmetrically and results in a net force in the direction of transport driven by binding to cytoplasmic dyneins (Athale et al., 2014). In contrast, in mouse oocyte meiosis a gradient of dynein but uniform MT lengths were seen to bias the positioning of MTOC asters (Khetan and Athale, 2016). However, the size scale of budding yeast cells is a few micrometers, compared to the diameter of vertebrate oocytes ranging between 10^2^ to 10^3^*µ*m. The fact that astral MT lengths and cell diameters are comparable in budding yeast, suggests frequent encounters of cMTs with the membrane. Thus, the polarized localization of dyneins at the membrane can be seen as a robust mechanism for generating a polarized force on astral MTs nucleated from SPBs.

**Table 4:** Quantification of experimental measures of SPB motility during the budding process for the two SPBs that enter the daughter-cell (D) and mother cell (M).

| *Motility measure* | *SPB-D* | *SPB-M* |
| --- | --- | --- |
| $D_{eff}$ | $1.6 \pm 4 \times 10^{-4} \mu\text{m}^2/\text{s}$ | $2.416 \pm 0.3 \times 10^{-4} \mu\text{m}^2/\text{s}$ |
| $\alpha$ | $1.96 \pm 1.89$ | $1.6 \pm 1.7$ |
| $c_1$ | $0.589 \pm 1.0983 \mu\text{m}^2$ | $0.141 \pm 0.179 \mu\text{m}^2$ |
| $D'_{eff}$ | $2 \pm 3.5 \times 10^{-5} \mu\text{m}^2/\text{s}$ | $2.5 \pm 3.6 \times 10^{-5} \mu\text{m}^2/\text{s}$ |
| $v_{drift}$ | $17 \pm 18 \text{ nm}/\text{min}$ | $4.1 \pm 8.4 \text{ nm}/\text{min}$ |
| $c_2$ | $0.5733 \pm 1 \mu\text{m}^2$ | $0.252 \pm 0.169 \mu\text{m}^2$ |
| $\chi$ | $1.16 \pm 0.55 \times 10^{-1}$ | $7 \pm 4 \times 10^{-2}$ |

**Figure 3:**
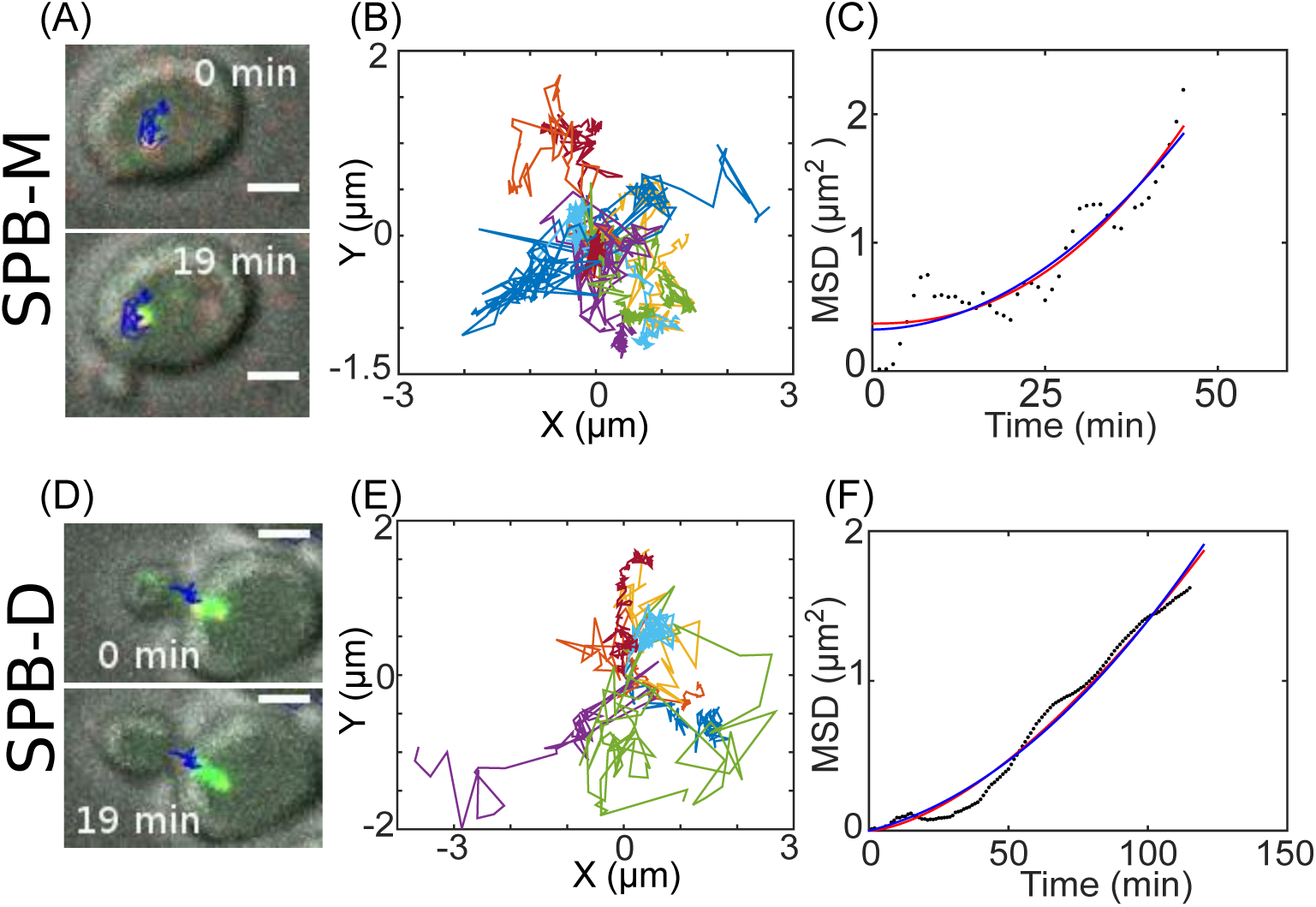
SPB mobility during mitosis. (A,D) Representative images of budding cells in DIC (gray) merged with the SPC42-3xmCherry (red) and GFP-Tub1 (green) channels are overlaid with the trajectories of the tracked SPBs entering the (A) mother cell (SPB-M) and (D) daughter cell (SPB-D). Scale bar 2 *µ*m. These images correspond to Videos SV3(a) and SV3(b). (B,E) The XY-trajectories of (B) SPB-M (n=20) and (E) SPB-D (n=16) are plotted. Colors represent individual trajectories. (C,F) Representative MSD profiles of SPBs from experiment are plotted with time (dots) for (C) SPB-M and (F) SPB-D and fit to an anomalous diffusion model (red, Equation 2) and diffusion with drift model (blue, Equation 3). Scale bar 2 *µ*m.

While the transport, binding and activity of dynein to the cortex have been quantified, it remains unclear if the quantified cluster sizes of dynein and the density in the bud-cell cortex are sufficient for both ‘search and capture’ of cMTs nucleated by the SPB-D and for force generation to pull the nucleus to the bud cortex.

### Model of ‘search and capture’ of cMTs by cortical dynein and estimating pulling forces

The model of the pre-budding stage of nuclear mobility due to diffusion and MT polymerization, was extended to account for the budding stage. We used a minimal 2D model representing the budded cell as an ellipse, with dynein motors localized in an arc reflecting the pre-anaphase polarized localization of dynein in the bud (Figure 4A). The two SPBs are embedded at diametrically opposite sides of the periphery of the nucleus, and as in experiment the SPB migrating towards the daughter cell is referred to as SPB-D and that which stays in the mother cell SPB-M. Each SPB nucleates two dynamic MTs with their plus-ends free and minus-ends embedded in the SPB. Length fluctuations of MTs are determined by dynamic instability. MT dynamics turnover is reported to be higher in pre-budding cells as compared to budded (Carminati and Stearns, 1997). We therefore choose dynamic instability parameters previously reported from pre-anaphase cells (Fees et al., 2017). In our model, the dynein motor is modeled as a stochastic stepper with force- and direction-dependent detachment rates, as described before (Jain et al., 2019). Unlike in a model of *Schizosaccharomyces pombe* nuclear positioning, we do not implement length dependent MT catastrophes (Foethke et al., 2009), in order to maintain the simplicity of the model. The spring-like motors immobilized at the cell cortex stochastically bind to, i.e. ‘capture’, cMTs if they are within the minimal binding distance and once bound, walk to the minus-end of the MTs, embedded in the SPB. The dynein pulling forces result in MT sliding and movement of the nucleus (Figure 4A,B). As expected increasing density of motors results in a greater degree of polarized motion of the SPB-D into the bud region, with the SPB-M following (Figure 4C).

**Figure 4:**
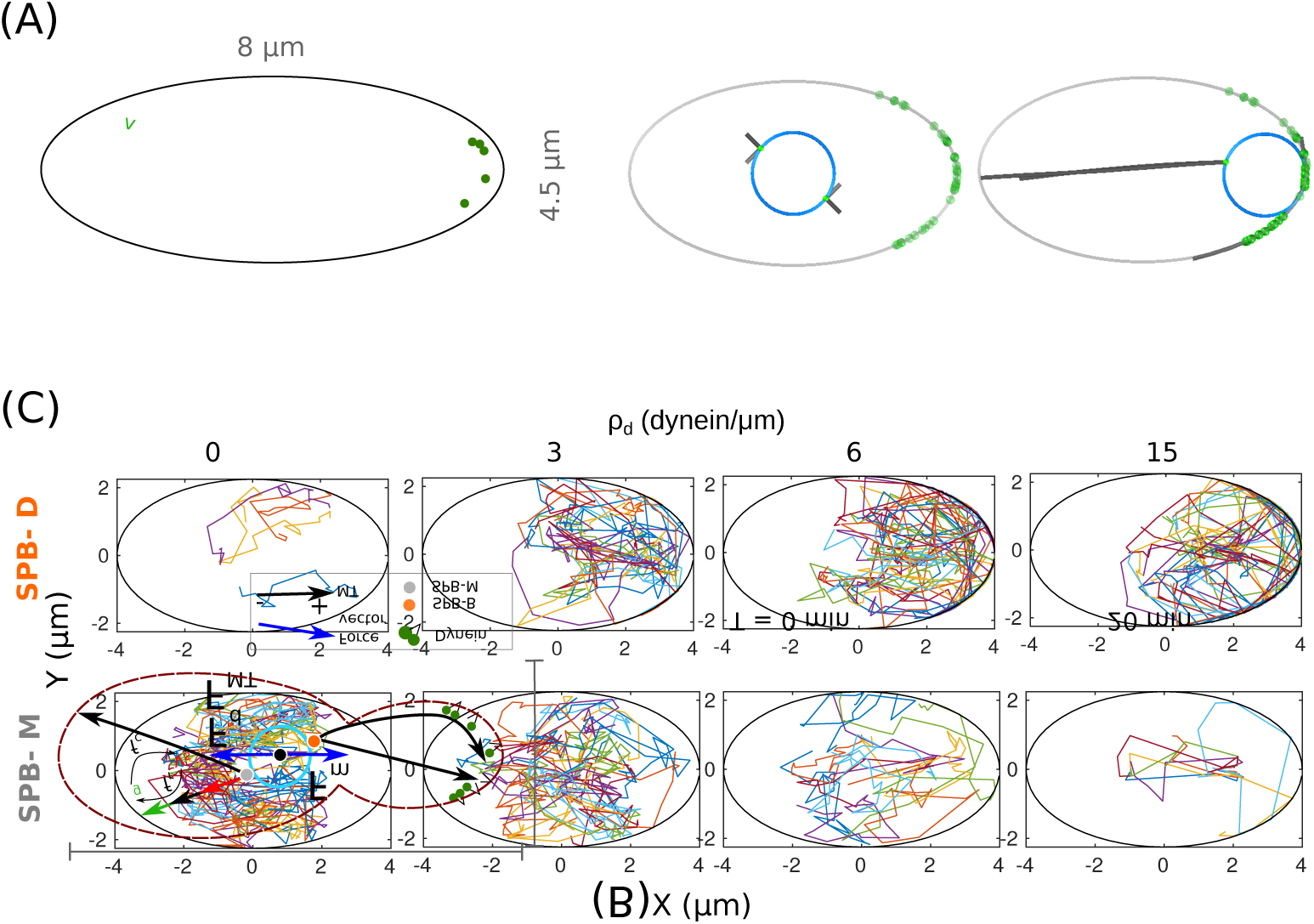
Simulating nuclear positioning by dynein localization in the bud. (A) Schematic of the geometry of a budding yeast cell represented as an ellipse with polarized motors (magenta) acting on dynamic MTs nucleated from the SPB-D (green) embedded in the nuclear envelope, like the diametrically opposite SPB-M (orange). Dynein is localized in an arc along the cortex. *θ*= 60 deg while *f*_*c*_, *f*_*r*_: frequencies of catastrophe and rescue; *v*_*g*_, *v*_*s*_: velocities of growth and shrinkage, respectively. (B) Representative simulations of polarized dynein from the start and end time points demonstrate the convergence of SPB-D to the bud tip. Motor density in the bud region is 6 motors/*µ*m. (C) XY trajectories of SPB-D and SPB-M are plotted for cortical dynein densities 0, 3, 6 and 15 motors/*µ*m. Colors indicate individual SPB tracks. Simulation time = 20 minutes, N_*runs*_ = 20.

In order to quantify the degree to which the polarized dynein pulling affects nuclear positioning, we have divided the simplified geometry of the elliptical model of the budding cell into 1/4th regions along the mother-bud axis represented by the long-axis of the ellipse (Figure 5A). The model predicts a saturation of the fraction of nuclei entering the 1/4th of the cell representing the bud with dynein density *ρ*_*d*_ higher than 12 motors/*µ*m (Figure 5B). The saturation for *ρ*_*d*_ ≥ 12 motors/*µ*m is also seen in terms of the time taken for a nucleus to first enters the 1/4th polar region, i.e. first passage time (Figure 5C) and maximal velocity and the inferred drag force (Figure 5D).

**Figure 5:**
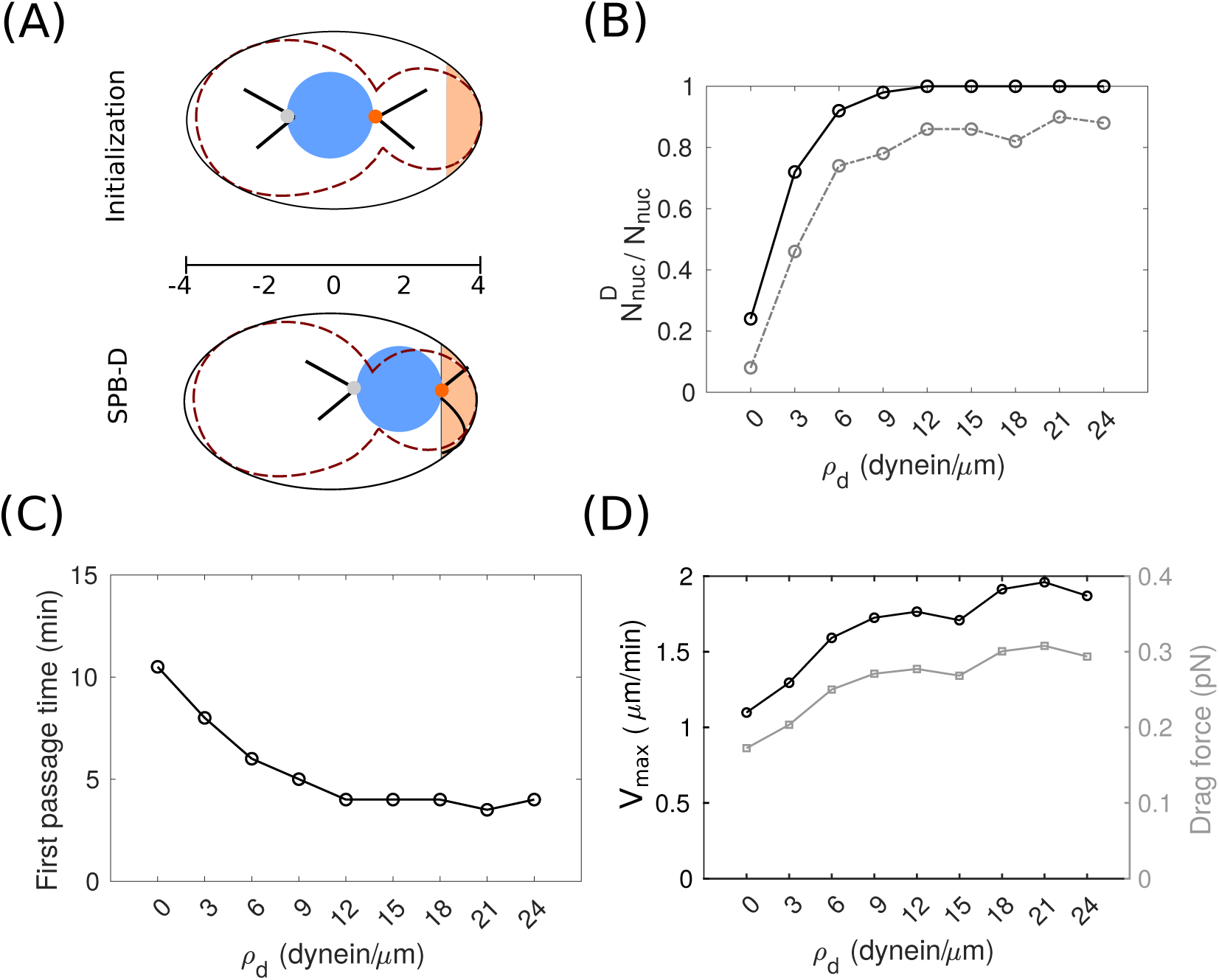
Nuclear convergence by ‘search and capture’ of cMTs and dynein pulling forces. (A) The schematic represents the geometry of the minimal model of a budding cell. The geometry of mother-daughter cell (dashed brown line) is simplified to an ellipse. The mother-daughter axis is divided into the polar 3/4th of the daughter cell (orange) and any part of the nucleus entering this region is considered to have been positioned at the bud-neck. (B-D) The effect of increasing dynein density (motors/*µ*m) in the bud-cell cortex evaluated in terms of (B) the fraction of total nuclei simulated, *N*_*nuc*_, whose boundary enters the polar region of the daughter cell, 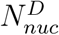 at the least once (solid line) or at the end of 20 min of simulations (dashed line), (C) the median time it takes the first nucleus to enter the polar-zone of the bud, the first passage time, and (D) the maximal velocity of nuclear movement and the corresponding effective drag force. All measures are averaged over 50 simulation runs for each case, *η*_*cyto*_ = 0.5 Pa s, *r*_*nuc*_ = 1 *µ*m, total time of simulation is 20 minutes.

Thus we find a minimal motor surface density of 12 motors*µ*m is required for reliable positioning of the nucleus. However, it does not say much about the number of motors engaged. In order to test the model predictions we will also require an experimental estimate of a similar number.

### Number of dyneins pulling nuclei based on experimental motility, force-balance and simulation

We proceed to use a force-balance approach to estimate the minimal number of motors required to transport the nucleus, from experimental data. The directional motility of the nucleus during budding is influenced by two broad classes of forces, (a) those that resist transport into the bud: *F*_*MT*_, due to MT pushing and polymerization at the membrane and *F*_*d*_, the drag force opposing the motion of the nucleus into the pole, and (b) those driving the nucleus into the bud: *F*_*m*_: motor pulling forces (Figure 4A). Therefore at any stage, the combination of forces acting on the nucleus can be termed as the net force, 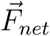, which we define as the vector sum of the forces, 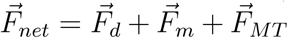 where 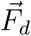 is the drag force opposing the motion, 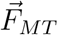 is the force of MT pushing by polymerization into the cell and 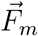 is the motor force pulling the nucleus to the pole. At equilibrium all forces cancel out and 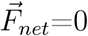. However, in the budding process, the directed motion of the nucleus suggests the dominance of the pulling force. In our simple model, it is only the force of cortical dynein. The forces opposing it are the drag force due to cytoplasmic viscosity and the force of MTs opposing the movement. We can simplify our expression for net force then as the difference in force magnitudes as:

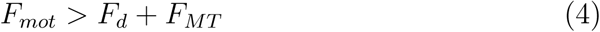

The movement of the nucleus and the interphase viscosity *η*_*cyto*_ allow us to estimate the drag force as:

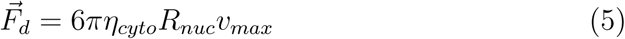

where *R*_*nuc*_ is the nuclear radius, *v*_*max*_ is the maximal velocity of transport. If we assume that the velocity of nuclear translocation driven by motors reaches a maximal value, measured in experiments to be 8.3 nm/s. We consider this to be an effective ‘terminal velocity’. Thus *F*_*d*_ is 0.039 pN and the pushing force of MTs *F*_*MT*_ depend on the number of MTs, 17 pN for one and 35 pN for two MTs acting simultaneously (Das et al., 2014). If we assume the total force exerted by the motors to be *F*_*m*_ = *N*_*m*_ *· f*_*s*_ where *f*_*s*_ is the single motor stall force and simplifying Equation 4 combined with the linear dependence of force on the number of motors *N*_*m*_, we can estimate the minimal number of motors *N*_*m*_(*min*) required as:

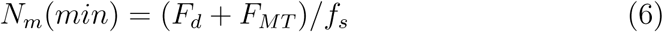

This is the threshold number of motors, above which the opposing forces acting on the nucleus can be overcome and the nucleus positioned. Given the single motor yeast cytoplasmic dynein stall force ranges between 4 to 7 pN (Reck-Peterson et al., 2006; Gennerich et al., 2007; Belyy et al., 2014; Cleary et al., 2014) and the maximal velocity of SPB-D in experiments, we arrive at *N*_*m*_(*min*) of 2 or 5 motors per nucleus engaged with either one or two cMTs (Figure 6A). This is consistent with the *in vivo* measurement from previous work of 3±1 dimers of cortical dynein molecules sliding MTs in the bud cell of *S. cerevisiae* based on Dyn1-3mCherry puncta quantification (Markus et al., 2011) and fluorescently labelled Num1 clusters that bind 1:1 to dynein seen to be interacting with astral MTs during capture and sliding (Omer et al., 2020).

**Figure 6:**
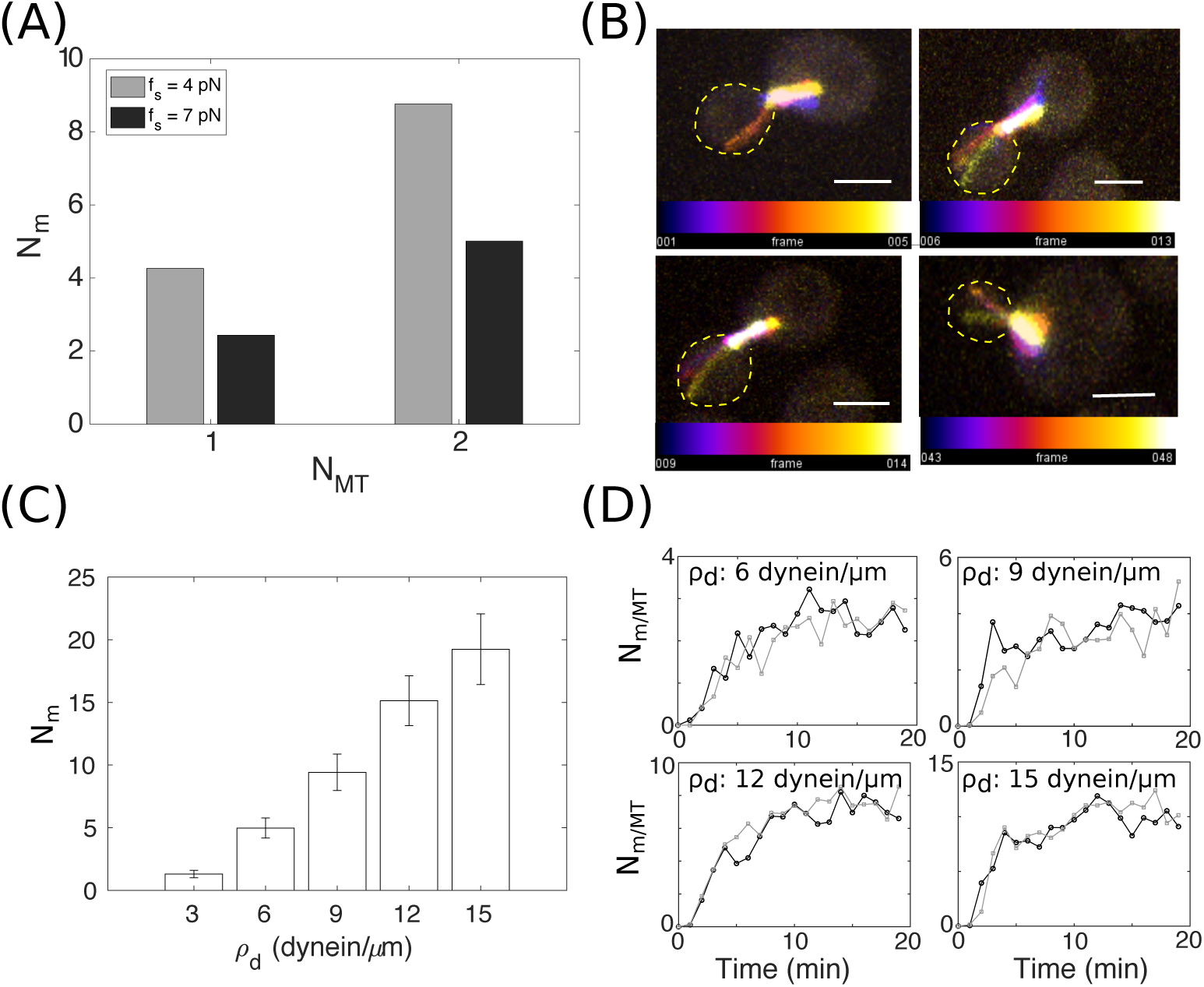
Number of dyneins pulling on MTs in the daughter cell. (A) The minimal number of motors per nucleus (*N*_*m*_) calculated based on force-balance required to drive nuclear positioning against MTs are plotted a function of number of opposing MTs (*N*_*MT*_) for dynein stall forces *f*_*s*_=4 (grey) and 7 pN (black). (B) Representative images of GFP-labelled MTs projected from time-series of 4 different budding cells. The background GFP signal allows the identification of the daughter cell outline (yellow dashed line). The colorbar indicates frame number. A frame was acquired every 1 minute. Scale bar: 2 *µ*m. (C,D) Simulation predictions of the (C) number of motors (*N*_*m*_) at the end of 20 min simulation bound to cMTs in the daughter cell (mean ± s.e.) and (D) average values of motors/MT engaged with the 2 astral MTs (black, grey) as they pull at the nucleus. Values plotted are means from 50 simulations for increasing cortical dynein density *ρ*_*d*_.

A projected time-series of cMTs in the bud suggests cMTs can spend 10-14 minutes in the bud near the cortex and encounters of the MT with the cortex appear to be either end-on or involve interactions of MT lengths ranging between 0.5 to 2.5 *µ*m (Figure 6B). It raises the question how such transient interactions that involve motors of a few nanometers in size can capture an MT of ≈ 25 nm diameter.

In simulations, we find 5 or more dyneins bound per nucleus on an average when the density dyneins is greater than 6 motors/*µ*m (Figure 6C). Indeed the kinetics of MTs binding to the cortex suggest an initial stage of capture of MTs by 1 motor, which rapidly increase and converge to a steady state, whose exact value depends on the motor density (Figure 6D). This is consistent with the idea of ‘reeling in’ of nuclei by cortically localized dyneins pulling on ‘captured’ cMTs. Indeed a quantitative comparison with experiments based on Num1 foci counting reveals 7.7 ± 3.6 foci per bud cell (Omer et al., 2018). Each Num1 cluster in turn has been seen to consist of 3 dimers associated 1:1 with 3 dynein dimers (Omer et al., 2020). This would amount to a surface density of ≈ 3.4 motors/*µ*m. While our simulation predicts a minimal motor density two-fold higher, the mean number of motors bound, and therefore force generated matches the experimental results of 4 (Markus et al., 2011) and 3 dimers of dynein (Omer et al., 2020). This difference in linear density between model and previous experimental reports could stem from dynein clustering, with each cluster consisting of 3 to 4 dyneins. Since each cluster contains sufficient motors to generate the minimal force required to overcome the opposing forces, it could explain how cells could use a lower average density of motors to generate the minimal force required. At the same time, this raises a separate question of the efficiency of ‘search and capture’ of cMTs by clusters of dynein as opposed to uniformly distributed motors. Future modeling in a realistic geometry with active transport and clustering of dynein could be used to address the physical basis of both correct positioning and correct orientation of nuclei in the budding process.

## Conclusions

In this study we use a model of nuclear mobility in pre-budding cells of *S. cerevisiae* driven by diffusion and MT pushing and by reconciling to experimental mobility measures of the SPB, estimate the effective cytoplasmic viscosity to be 0.5 Pa s. By tracking SPB-D mobility in the budding stage from experiments, we estimate minimal force that dynein motors must collectively exert in order to to position the nucleus, to overcome MT pushing forces and viscous drag. Our simulations of nuclear positioning by ‘search and capture’ and pulling of cMT demonstrate a cortical density of at least 6 dynein dimers per *µ*m is required to robustly ‘capture’ cMTs and position the nuclei to the bud neck while a minimum of 6 dyneins per nucleus are required to generate sufficient force. The model predictions compared to experiments suggest a functional role for motor clustering in dynein collective force generation once MTs are captured.

## Author Contributions

KJ designed and performed the experimental data analysis; SP performed the *S. cerevisiae in vivo* time-lapse experiments; NK performed the simulations; CAA designed the research wrote the manuscript and obtained funding.

## Acknowledgments

We would like to thank Gislene Pereira for the gift of GFP-Tubulin, SPB-mCherry *S. cerevisiae* strain and Mohan Balasubramanian for the support for using the microscope facility at the University of Warwick, U.K. A grant from the Dept. of Biotechnology, Govt. of India (BT/PR6715/GBD/27/463/2012) and IISER Pune core-funding to CAA supported the work. KJ was supported by a fellowship from DST-INSPIRE (IF-130394) and a research assistantship from SERB, Govt. of India (EMR/2016/005481). NK was supported by fellowships from CSIR (09/936(0128)/2015-EMR-1) and the Dept. of Biotechnology, Govt. of India (BT/PR16S91/BID/7/673/2016). SP was supported by Wellcome Trust (WT101885MA).

## 4 Supporting figures

**Figure S1:**
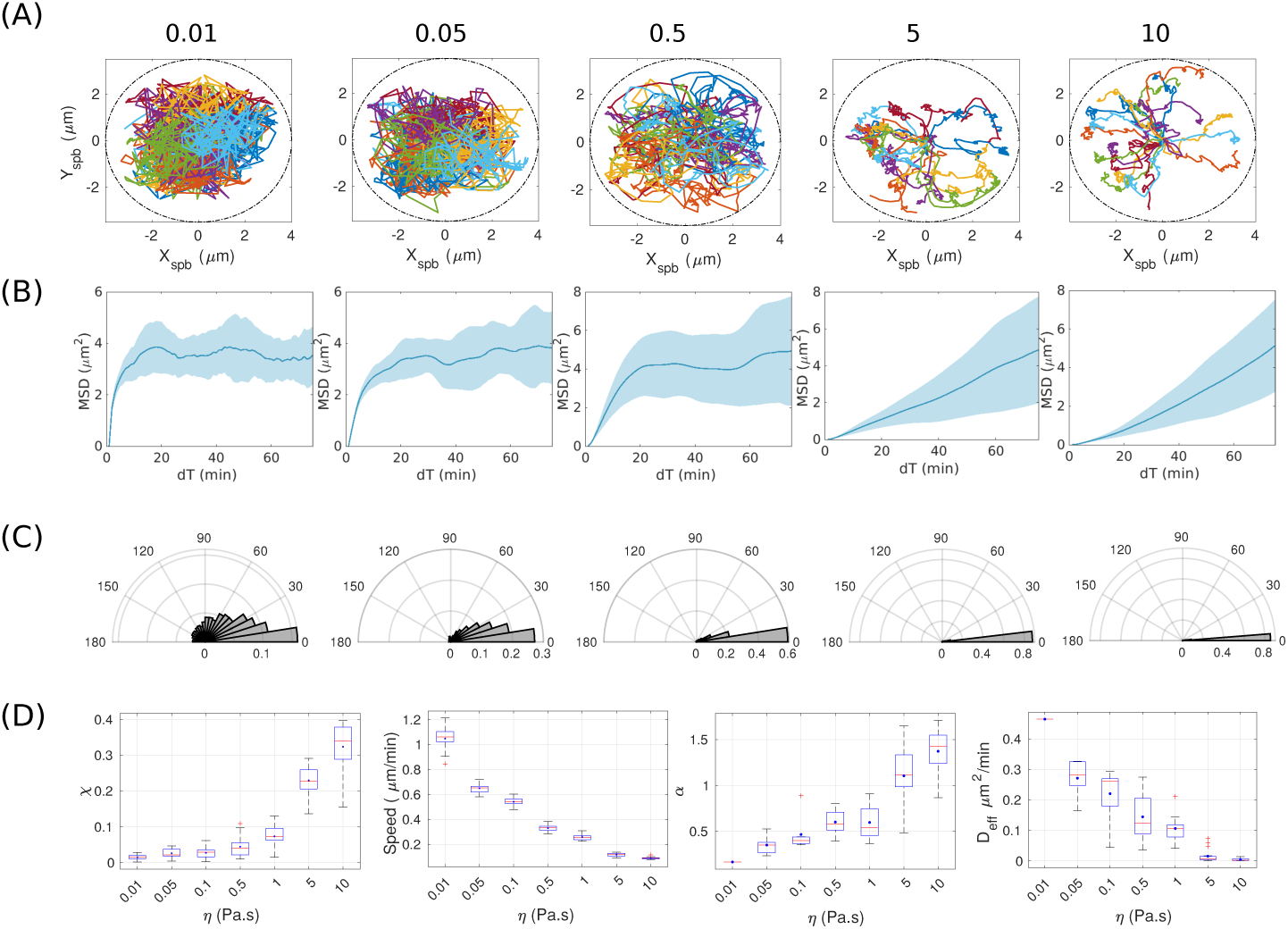
Effect of viscosity on mobility parameters of the nucleus in simulation. (A) The XY trajectories of SPB motility across varying effective cytoplasmic viscosity *η* were used to calculate (B) MSD with time averaged over particles (blue line) with the area indicating s.d. (C) The frequency distribution of the smallest angle between subsequent SPB positions *θ* indicates rotation of the nucleus. (D) The box plots represents measures of motility as a function of *η* for directionality *χ*, instantaneous velocity and fit parameters of the MSD fit to the anomalous diffusion model (Equation 2), the effective diffusion coefficient *D*_*eff*_ (*µ*m^2^/min) and anomaly parameter (*α*). Blue circles represent mean and red lines represent median values. N_*runs*_ = 50.

## 5 Supporting data

Data S1: **Experimental statistics of pre-budding SPBs**. The averaged statistics of trajectories obtained from single particle tracking (SPT) of SPB mobility in pre-budding cells are summarized in this table.

Data S2: **Calculations for force-balance of nuclear migration from experiment and simulation**. The force-balance calculations to estimate the minimal number of dynein motors (*N*_*m*_) are described in this MATLAB file.

### 5.1 Supporting videos

**Video SV1:**
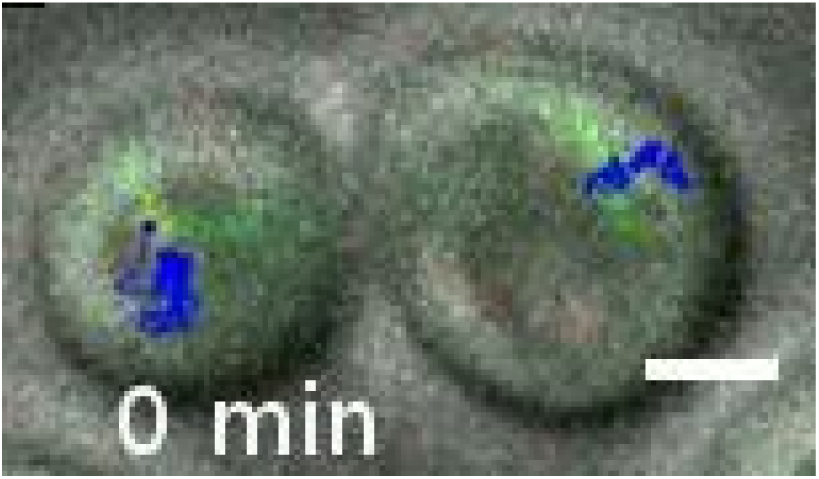
Diffusive SPB mobility. A representative movie of a time series of two neighboring yeast cells in the pre-budding stag, selected based on morphology in DIC (gray). The Spc42-3xmCherry (red) and GFP-Tub1 (green) channels are overlaid and the SPB mobility tracked (blue). Scale bar: 5 *µ*m, time: minutes.

**Video SV2:**
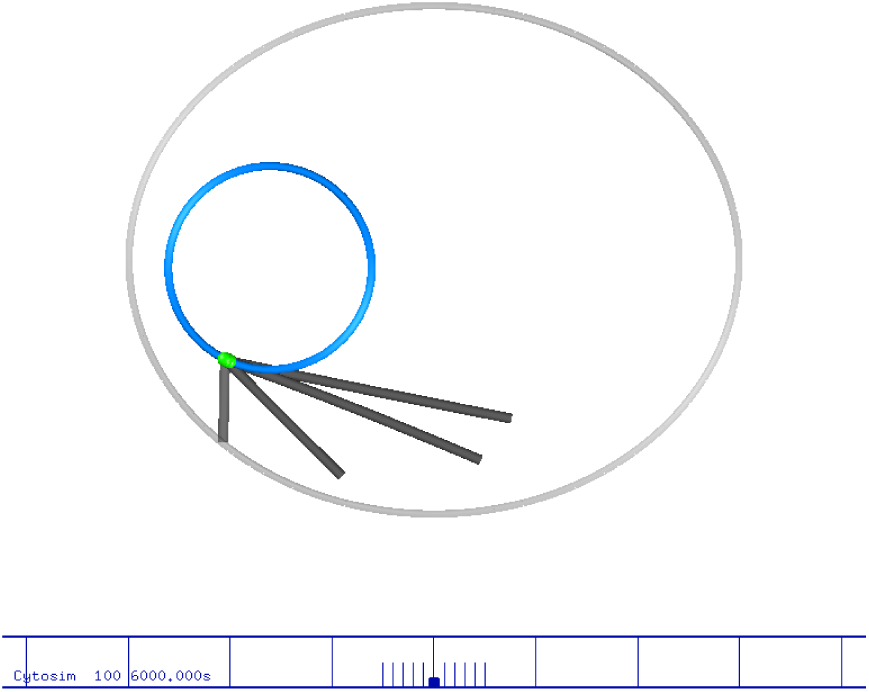
Simulation of diffusive mobility of SPB attached to a nucleus. The output from a simulation run of a nucleus (blue) with a pair of SPBs anchored close to each other in the membrane (green), each nucleating a pair of astral MTs. MT dynamic instability drives filament length fluctuations, while pushing at the membrane and diffusion position the nucleus. Viscosity 0.5 Pa.s, total simulation time 20 min, Δt=10 s, cell radii are 3 *µ*m (major-axis) and 2.5 *µ*m (minor-axis), radius of the nucleus 1 *µ*m. Each block of the scalebar corresponds to 1 *µ*m. Time in seconds.

**Video SV3:**
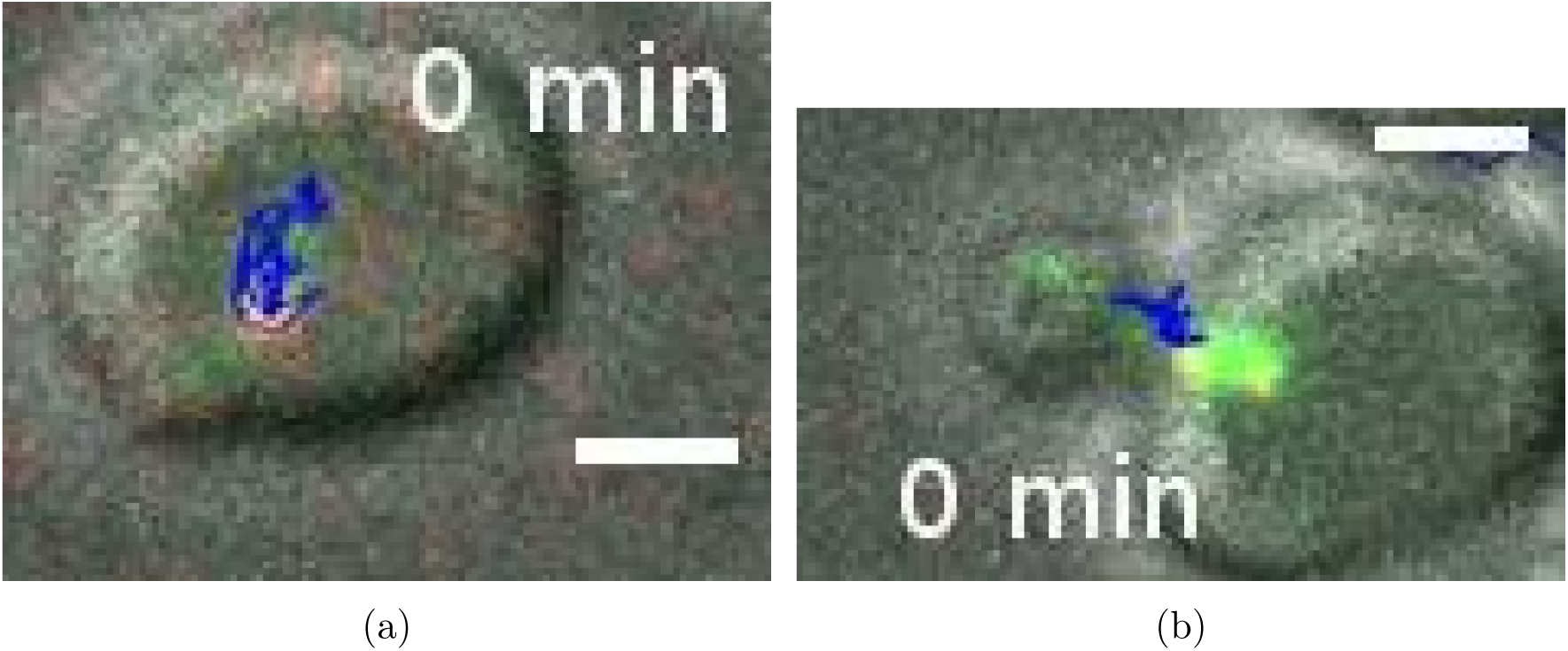
SPB mobility in the budding stage. (a),(b) Representative movies of time series of SPB mobility during budding where the tracks of either the (a) SPB that remains in the mother cell or (b) that enters the bud cell are seen. The cell morphology is seen in DIC (gray), while Spc42-3xmCherry labels the SPB (red) and GFP-Tub1 labels MTs (green). All channels are merged and overlaid with the tracks of the SPB (blue). Scale bar: 5 *µ*m, time: minutes.

**Video SV4:**
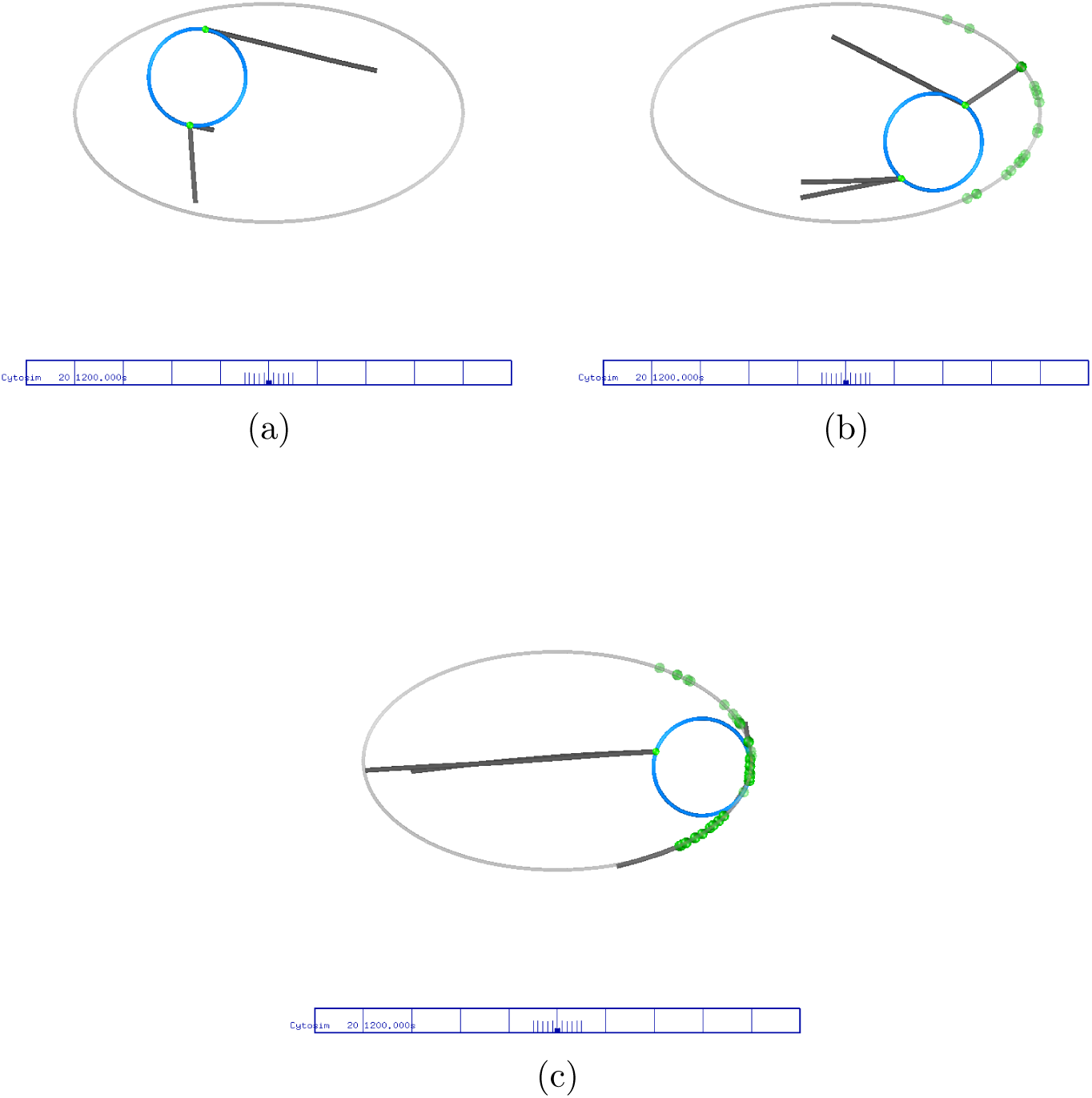
Simulating nuclear ‘search and capture’ by cortical dynein. (a)-(f) Simulations of the simulated yeast nucleus (blue) with a pair of SPBs at diametrically opposite ends of the nucleus with 20 minutes represents the position of the nucleus for varying densities of dynein at the cortex. *ρ*_*d*_ densities are (a) 0,(b) 3 and (c) 6 motors/*µ*m. Scalebar corresponds to 10 *µ*m.

